# Protein profiling in cancer cell lines and tumor tissue using reverse phase protein arrays

**DOI:** 10.1101/144535

**Authors:** Xiaohong Jing, Weiqing Wang, Nicholas P. Gauthier, Poorvi Kaushik, Alex Root, Richard R. Stein, Anil Korkut, Chris Sander

## Abstract

Reverse phase protein array (RPPA) technology is an antibody-based high-throughput assay for protein profiling of biological specimens that allows for many measurements with very small amounts of cell lysate. Here, we report the sensitivity, reproducibility, and accuracy of a particular RPPA platform called Zeptosens. We customized the RPPA protocol for our in-house setup, and measured more than 80 total protein and phospho-protein levels in various cancer samples, including cell lines, organoids, tumor chunks, core needle biopsies, and laser-capture microdissected tissue samples. We discuss pros and cons of the RPPA platform, and describe results from profiling 15 cancer cell line cells using RPPA.

## Introduction

Reverse phase protein array (RPPA) technology is an antibody-based assay that allows rapid measurement of protein expression levels in a large number of biological samples. One of the major advantages of this technology is that it can semi-quantitatively measure many low abundance analytes, such as phosphorylated signaling proteins, from small amounts of cell lysate^1–3^.

Protein microarray formats can be divided into two major classes: forward phase arrays and reverse phase arrays. In forward phase protein arrays, each spot contains one type of immobilized antibody or bait protein such that each array is comprised of hundreds of antibodies. Each array is incubated with one biological sample, such as a cellular lysate, and multiple analytes from that sample are measured simultaneously. In contrast, the RPPA format immobilizes an individual biological sample in each array spot such that an array is comprised of hundreds of different samples. Each array is incubated with one detection protein (i.e., antibody) that is directly compared across multiple samples^4^. Both RPPA and forward phase arrays allow hundreds of samples to be probed by hundreds of antibodies. If sample volumes are small, and if there are a large number of samples compared to antibodies, RPPA is the preferred method.

Cellular physiology and disease are regulated by protein signaling pathways. The activity of these pathways can be monitored through changes in protein levels and post-translational modifications, such as phosphorylation. RPPA provides a high-throughput and sensitive tool to profile these pathways, which can help identify new biomarkers and potential therapeutic targets in laboratory research and clinical trials^1–6^. The quantitative and rich proteomic information generated by the RPPA technology enables us to go beyond interrogation of individual protein activities to comprehensively analyze changes in signaling. When combined with statistical inference algorithms, it also provides the opportunity to build comprehensive and predictive computational models of the response of cellular systems to perturbations^7–8^.

Here we report a series of experiments using the commercially available Zeptosens Technology RPPA platform (Bayer Technology Services, no longer sold as of December 2016)^9,10^. We set up the platform for accurate and reproducible measurements of protein expression from cultured cells and diverse tumor samples.

## The Zeptosens RPPA protocol

Below is a description of our customized Zeptosens RPPA protocol, which can also guide workflow development for other RPPA platforms.

### Required equipment and materials

- t-PREP Biomarker Extraction System (Covaris, #500305, 520097, 500312)
- CLB1 and CLB96 lysis buffer, CSBL1 spotting buffer, CAB1 buffer (Bayer Technology Services)
- EZQ Protein Quantitation kit (LifeTechnologies, #R-33200)
- Pierce Coomassie Plus (Bradford) Assay Kit (LifeTechnologies, #23236)
- Biomek FXP Laboratory Automation Workstation (Beckman Coulter)
- Nano-Plotter NP2.1 automatic pipetting system (GeSiM)
- ZeptoFog station (Bayer Technology Services)
- ZeptoChip (tantalum pentoxide-coated glass chips, Bayer Technology Services)
- SYPRO Ruby Protein Blot Stain (Life Technologies, #S-11791)
- ZeptoREADER microarray fluorescence readout system using Planar Wave Guide technology (Bayer Technology Services)
- ZeptoView 3.1 image analysis software (Bayer Technology Services)

#### 1. Sample preparation

Cell lysates are prepared from either cultured cancer cell lines, organoids, or primary patient tissue in CLB1 urea-based lysis buffer^11^. CLB96 buffer is used for cells cultured in 96-well plates. For tissue homogenization, we tested the FastPrep bead mill and Covaris t-PREP systems, and obtained a higher yield of extracted protein lysate with Covaris t-PREP. Phospho-protein states should be better preserved with Covaris t-PREP since samples remain frozen while pulverized. We therefore chose to use this system going forward.

We aim to collect 40 μL of cell lysate in CLB1 buffer at 2 mg/mL, or 60 μL of cell lysate in CLB96 buffer at 0.2 mg/mL. We are able to obtain this amount of lysate from individual wells of a 6-well plate and from core needle biopsy tissue samples, and it is sufficient for both chip spotting and protein concentration measurements before spotting. If less lysate is obtained (e.g., total protein concentrations as low as 1 mg/mL or volumes as low as 10 μL), the RPPA experiments are still doable.

#### 2. Quantitation of protein concentrations and sample dilution

We measure total protein concentrations for two purposes: 1) to ensure that similar concentrations of all samples are loaded on the arrays, and 2) for protein loading normalization during data analysis. We tested three protein quantification methods. The Pierce Coomassie Plus (Bradford) Assay Kit and EZQ Protein Quantitation Kit are used before diluting and spotting, while SYPRO Ruby Protein Blot on-chip staining is used after spotting to determine the relative total protein levels and normalize for loading differences. The EZQ kit demonstrated a larger and more suitable dynamic range (0.04-5 mg/mL) than the Coomassie Plus Bradford assay (1-12 mg/mL).

Ideally, protein concentrations for all samples are manually adjusted to 2 mg/mL before spotting. Samples at a lower concentration are used as is. Samples are then further diluted 10X with CSBL1 spotting buffer to generate a working concentration of approximately 0.2 mg/mL. We then prepare a linear dilution series of the samples (75%, 50%, and 25% for final concentrations of 0.2, 0.15, 0.1, and 0.05 mg/mL) to increase confidence in quantitation and verify that our measurements fall within the dynamic range. The dilutions are prepared by the Biomek FXP Laboratory Automation Workstation automatic pipetting system and are transferred to 384-well plates (20 μL/well) for array spotting. As suggested by Bayer Technology Services, these plates can be stored at -80 °C for several months before spotting.

#### 3. Array spotting

Diluted samples are printed onto tantalum pentoxide-coated glass chips (ZeptoChips) under environmentally controlled conditions (constant 53% humidity and 14 °C) using the non-contact printer Nano-Plotter NP2.1 from GeSiM. We optimize the spotting parameters (e.g., distance from pin to chip and setting piezo parameters) to achieve uniformly dispensed, round-shaped, well-aligned spots with no satellite spots. These parameters must be tested and adjusted for with each instrument, and we suggest following the general GeSim recommendations. To spot the chips, the piezoelectric tip of the Nano-Plotter aspirates the desired volume of each sample from the appropriate well of the 384-well plate, and then deposits 400 pL droplets onto arrays with one droplet per array. The chip and array layout is shown in Figure 1. There are six arrays on each chip and each array is stained with one antibody. Since the spotted volume is low, one can theoretically make thousands of antibody measurements from a single, small sample. A reference grid of Alexa Fluor-647 conjugated BSA is spotted onto each array.

**Figure 1:**
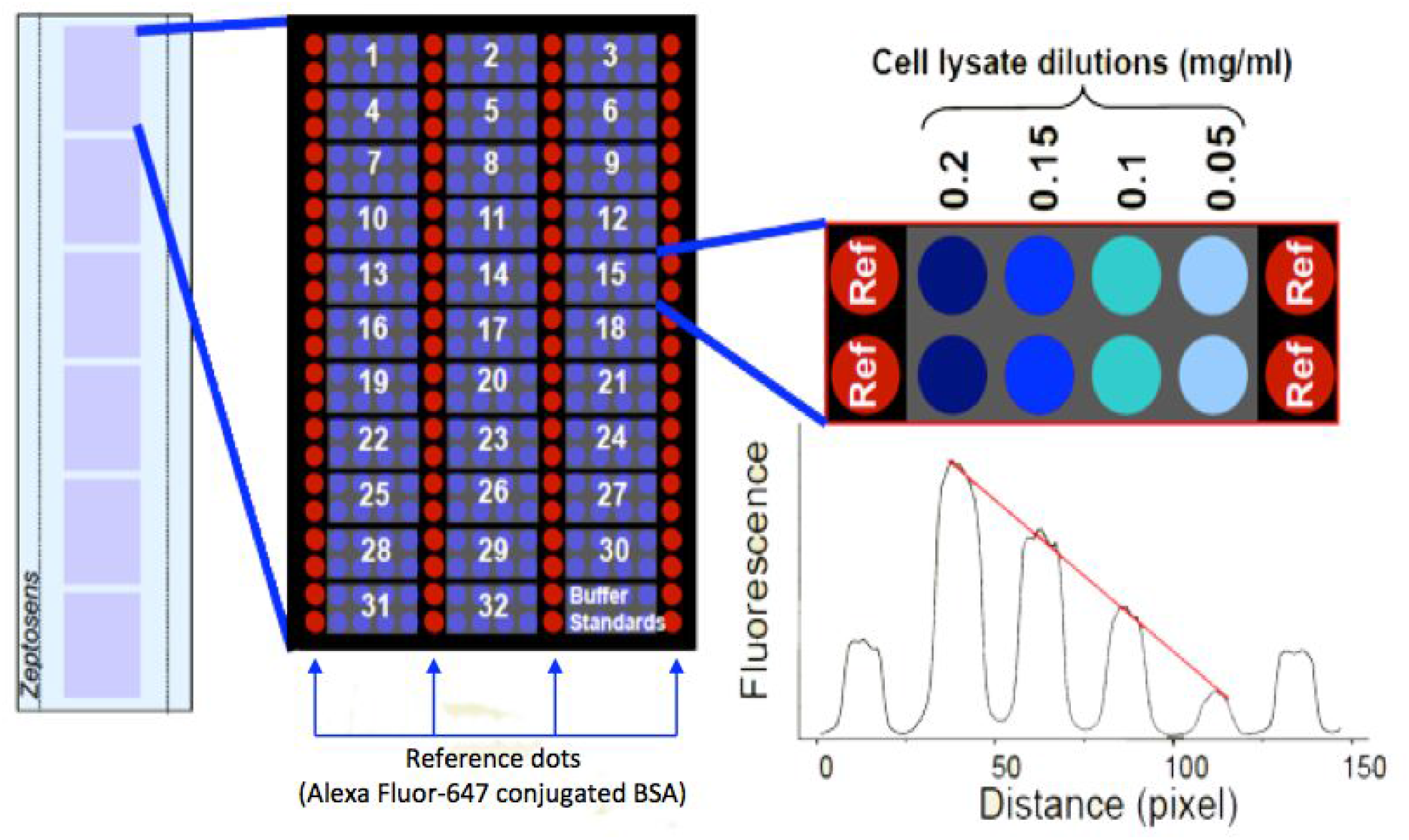
Zeptosens chip and array layout. There are six arrays on each chip and each array is stained with one antibody. A reference grid of Alexa Fluor-647 conjugated BSA is spotted onto each array. Each sample dilution series is spotted in between reference columns. Either 32 or 64 individual samples can be spotted onto each array. For a 32-sample layout, technical replicates are included in successive rows. Figure adapted from Zeptosens informational material provided by Bayer Technology Services.

#### 4. Blocking

After array spotting, the chips are blocked with an aerosol BSA solution using the ZeptoFOG blocking station (a custom designed nebulizer device) for 20 minutes. For SYPRO Ruby on-chip protein staining, the chips are blocked with non-protein based surface passivation reagents containing Tween-80. It is important that the ZeptoFOG blocking station is thoroughly cleaned because contamination from residual BSA can lead to decreased quality. The chips are subsequently washed in double-distilled H_2_O and dried. Blocked chips can be stored at 4 °C for a few months before carrying out the immunoassay (see Supplemental Data 1 for a comparison of chips stained with various antibodies immediately after spotting and chips stored for 2 months at 4 °C before staining).

#### 5. Antibody staining

Up to six spotted ZeptoChips can be inserted into one ZeptoCARRIER. Within the carrier, each chip is covered by a microfluidic structure forming a linear row of six flow cells (one flow cell per array). This setup enables a significant material cost savings compared to Western blots as only a minimal amount of antibodies and reagents are needed. For this immunoassay, we incubate the chips with primary antibodies for 20-24 hours followed by a wash with CAB1 buffer and a 2.5 hr incubation with Alexa Fluor-647 conjugated secondary antibody detection reagents. Antibodies are diluted in CAB1 buffer. After a final wash step in CAB1 buffer, the chips are ready for imaging.

#### 6. Fluorescence signal measuring

The immunostained chips are imaged using the ZeptoREADER instrument, which utilizes planar waveguide technology^12^. As reported by Bayer Technology Services, this technology increases the detection sensitivity up to 50-fold over conventional fluorescence. Two different fluorescence channels can be used: red (excitation 635 nm, emission 675 nm centered, width 50 nm) and green (excitation 532 nm, emission 572 nm centered, width 50 nm). For maximal throughput, 10 carriers can be stacked inside the reader, and each carrier can load up to six chips. For each array, four separate images are acquired using different exposure times ranging from 1 to 10 sec. Array images representing the longest exposure without saturation of fluorescent signal detection are automatically selected for analysis using the ZeptoView 3.1 software.

#### 7. Data analysis

Using the ZeptoView 3.1 software, a weighted linear fit through the dilution series is used to calculate the referenced sample fluorescence intensity (RFI) values for each sample replicate. Global and local normalization of sample signal to the reference BSA grid is used to compensate for intra-array spatial variation. Total protein concentrations as determined in Step 2 are used to normalize the sample loading amount for each sample. In practice, we perform protein loading normalization by dividing each RFI value by the estimated protein concentration (normalized RFI). Note that this normalization procedure is inappropriate if the linear fit of the dilution series generated from the four spots for an individual sample has a non-zero intercept, which can be verified from the raw fluorescence intensity data. The ZeptoView analysis software provides statistical methods to identify bad spots and remove them from further analysis. It is important during post-analysis to select the same exposure time for all arrays that have been stained with the same antibody. Choosing uniform exposure time is a prerequisite to make data from multiple arrays comparable. In practice, we select the median exposure of all ZeptoView-selected exposure times for each array. Further data analysis such as clustering, correlation analysis, and network modeling are carried out on these normalized data. Researchers at the MD Anderson Cancer Center have developed a web platform for visualizing and analyzing RPPA data called The Cancer Proteome Atlas (TCPA)^13^.

## Data reproducibility

We assessed the technical reproducibility of the Zeptosens RPPA method within and across arrays (intra-array and inter-array variations, respectively). We measured three analytes (AKT-pS473, PTEN, and Prohibitin) using lysates obtained from 15 cancer cell lines growing normally in culture. By choosing 15 lines, we aimed to assess a set of contexts with variable protein concentrations. These cell lines include five different cancer types: melanoma (SkMel-133, SkMel-173, SkMel-203, SkMel-207, SkMel-36), ovarian cancer (Caov-3, EFO-21, Ovcar3, Ovcar4, Skov3), liposarcoma (DDLS8817, LS141), breast cancer (MCF-7), and prostate cancer (LNCaP and LNCaP-AR). We also created a standard control lysate by mixing all 15 of the individual lysates.

### Intra-array variation

A dilution series with 4 concentrations of the 16 lysates (each cell line plus the standard control) was spotted in quadruplicate onto a single array. Technical replicates were highly correlated within the same array (Figure 2). Due to this reproducibility, we often do not print replicates on the same array and use the additional space for more samples.

**Figure 2:**
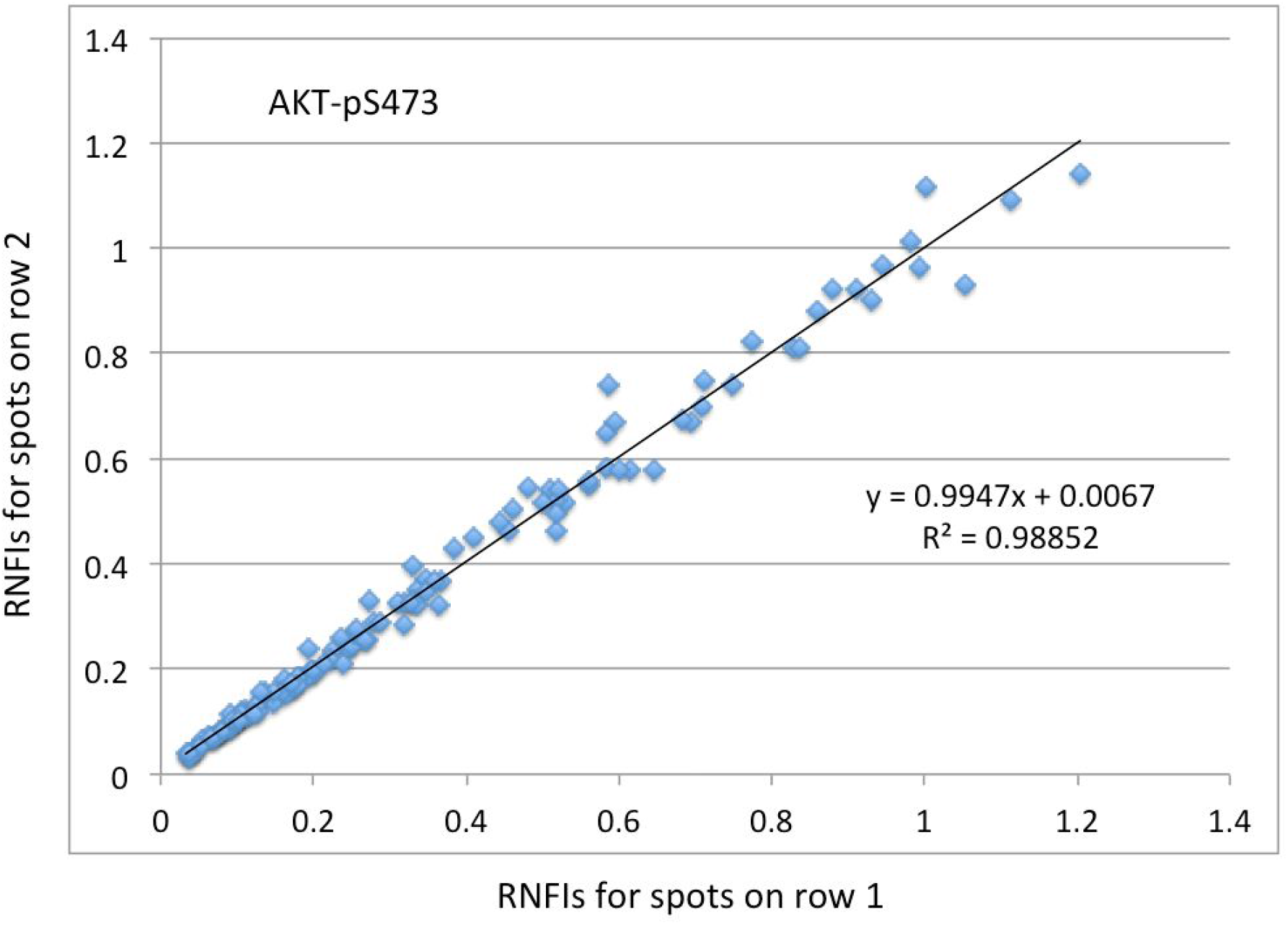
Intra-array correlation. Four dilutions of 16 samples were spotted in quadruplicate, stained with AKT-pS473, and measured with the Zeptosens RPPA platform. For each aspirate, the plotter dispensed two spots in successive rows (replicates). The referenced net fluorescence intensity (RNFI) is the intensity value for each spot normalized to the reference spots. Each datum depicts the agreement between the two replicate spots dispensed from the same aspirate onto successive rows.

### Inter-array variation

Using the same chip layout, four arrays were spotted with the 16 samples in quadruplicate and were stained with either PTEN or AKT-pS473 antibodies and imaged, and RFI values were calculated using the ZeptoView 3.1 software (Figure 3). Comparison of normalized RFI values (see Step 7) from these samples revealed that technical replicates were highly correlated across different arrays and that inter-array variation is not correlated with array location. The extent of correlation appeared to be antibody dependent. Pearson’s correlation coefficients for antibodies targeting PTEN, Prohibitin (data not shown), and AKT-pS473 were 0.994± 0.002, 0.950± 0.012, and 0.909±0.048, respectively. In general, it is not necessary to print technical replicates since intra-array and inter-array reproducibility are high. However, biological replicates are necessary to capture biological variability.

**Figure 3:**
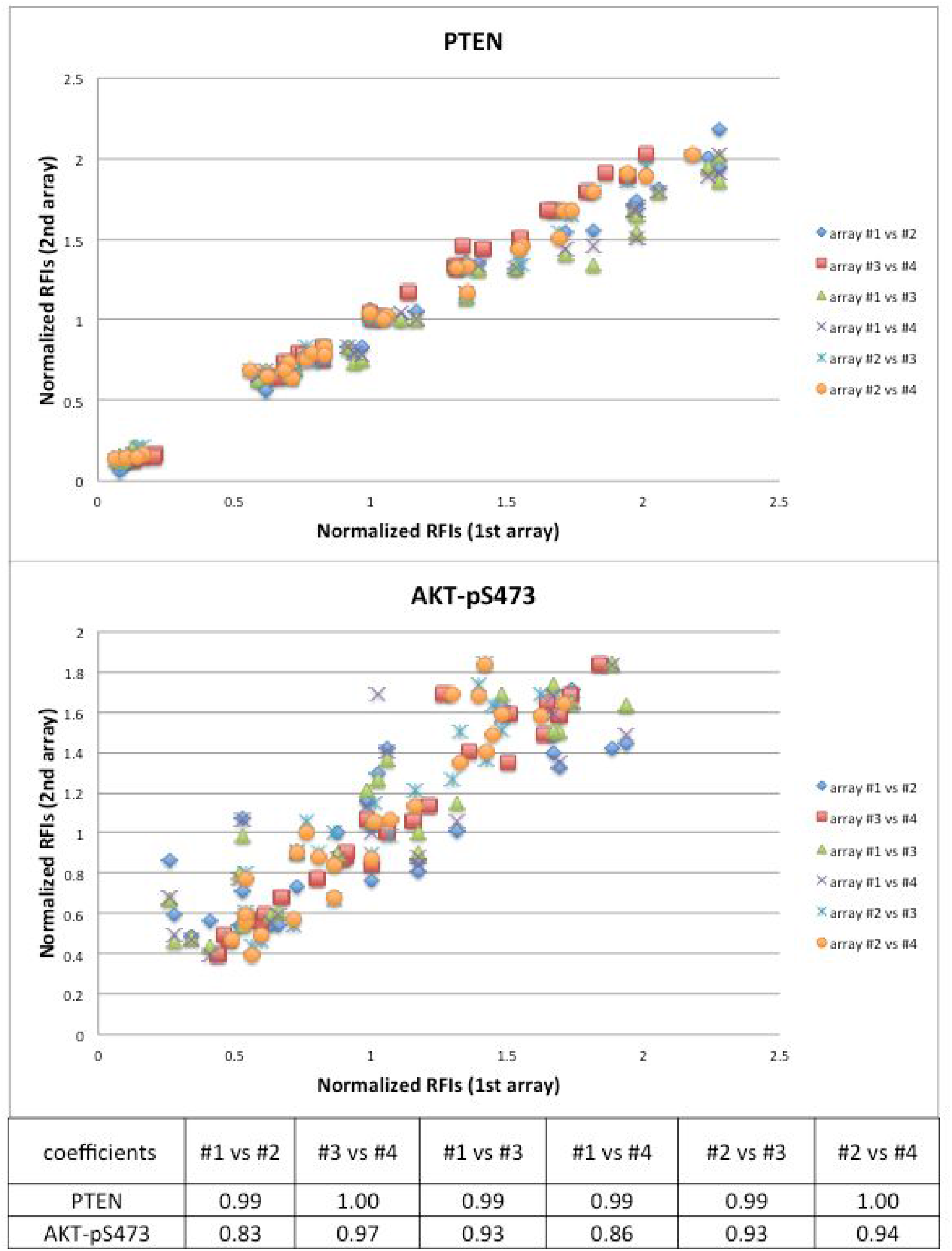
Inter-array correlation. Four replicate arrays (array #1 & #2 on one chip and array #3 & #4 on another) were spotted with the 16 samples in quadruplicate and were stained with the same antibody. RFI values were calculated for each sample by ZeptoView 3.1 software, and data were normalized for each array by subtracting the median RFI from all samples. Pearson’s correlation was determined across all possible array comparisons (table).

## Antibody validation and performance analysis

Antibody quality is critical for reliable, quantitative, and reproducible results with RPPA assays. Antibodies must have high specificity toward their target proteins. General approaches for antibody validation were described in the RPPA 2014 workshop report^1^. We consider an antibody to be specific if on a Western blot (1) there is a single band at the correct size or (2) there are multiple bands that are well characterized (e.g., different isoforms). High correlation between RPPA and Western blot can further increase confidence in particular antibodies. Bayer Technology Services has provided a list of ~300 validated antibodies, and we used 80 of these antibodies successfully in our RPPA runs. In addition, the functional proteomics RPPA core facility at MD Anderson Cancer Center has provided a list of more than 300 validated antibodies^14^. In general, the antibodies we have tested generate reliable and informative data (see Figure 4 and Supplemental Data 2).

**Figure 4:**
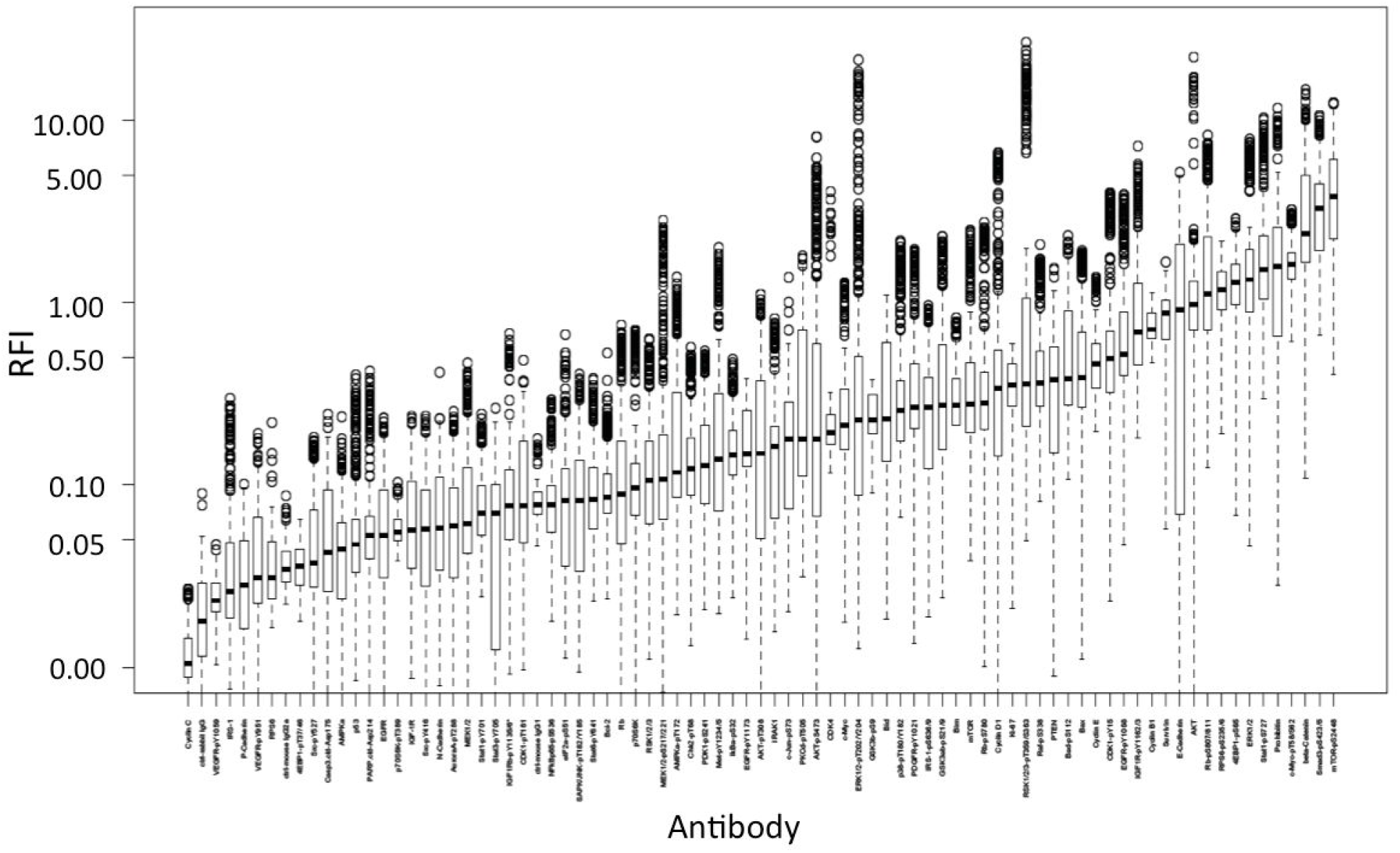
Antibody performance. Boxplot of RFI values acquired from 138,240 data points in our RPPA runs with lysates from cell lines. The RFI values were overall informative with average RFIs higher than 0.05. For most antibodies, we consider background noise to have an RFI around 0.01. See Supplemental Data 2 for the antibody names, as well as an overview of Zeptosens quality calls and RFI values for each antibody.

## Comparison of Zeptosens RPPA with Western blot

We performed RPPA and Western blot analysis with lysates prepared from the same 14 cell lines as described above and stained with four antibodies (pAKT, pERK, p-c-Jun, and PTEN). Note that 14 were chosen rather than all 15 in order to fit onto a single Western blot gel. Results from Western blots were consistent with those from RPPA runs (see Figure 5 and Supplemental Figure 1). In some cases, RPPA had a larger dynamic range than Western blot (e.g., for pERK).

**Figure 5.**
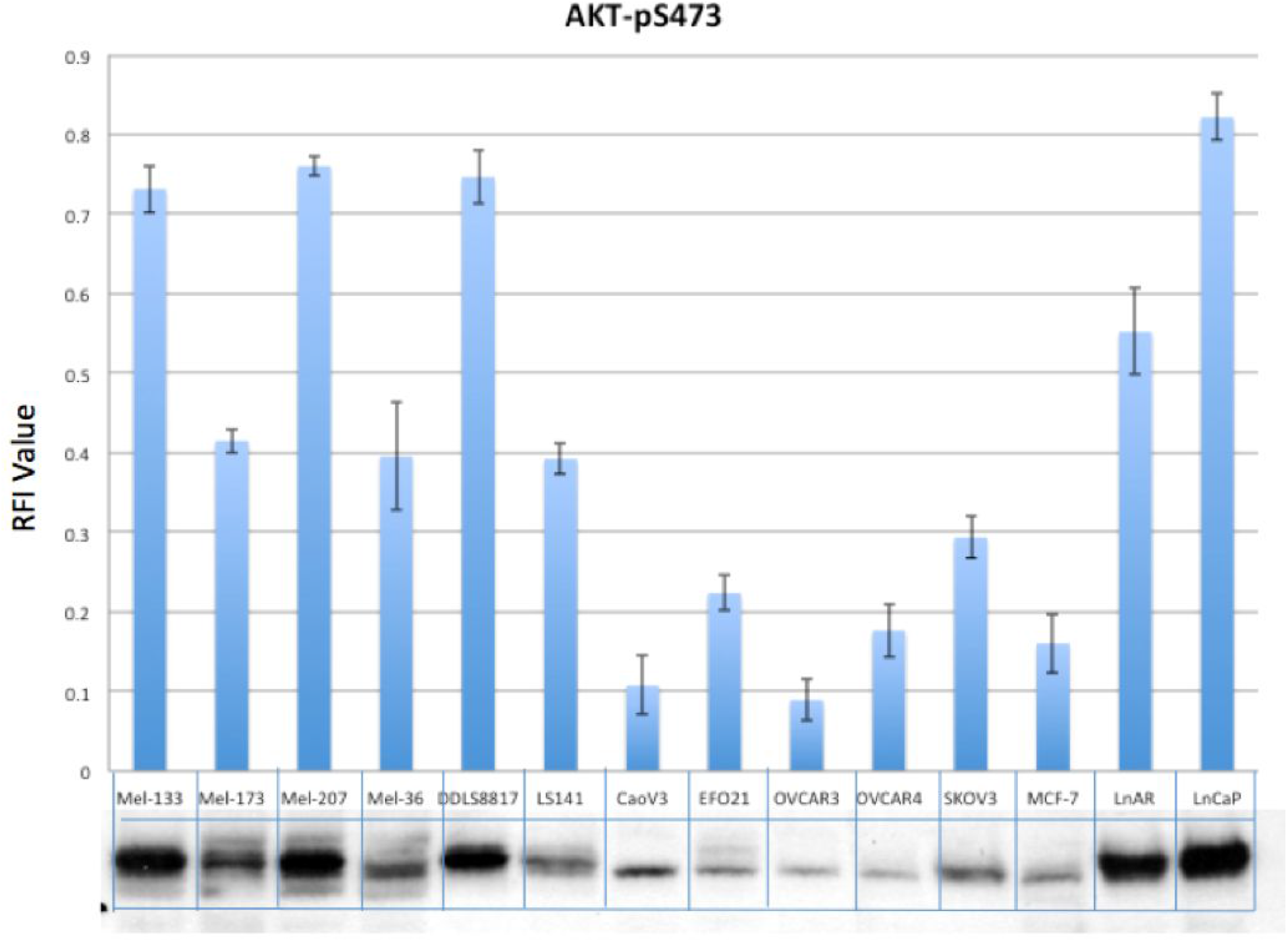
Comparison of RPPA and Western blot. Lysates for 14 cell lines grown in standard complete media were analyzed by both Zeptosens RPPA and Western blot. The Zeptosens RFI values (top, bar chart) and the raw Western blot images are shown for AKT-p473. See Supplemental Figure 1 for the additional three antibodies, as well as a comparison of Zeptosens RFI values to ImageJ-quantified Western blot results.

We estimate the overall cost for the Zeptosens RPPA to be ~$200 per chip, including buffers and antibodies. Therefore, the cost per data point is approximately $0.25 for Zeptosens RPPA (per RNFI value), whereas it is approximately $1-1.5 for Western blot. Note that our RPPA array layout utilizes a dilution series for every sample/data point (so the cost is $1 per RFI), whereas a Western blot typically utilizes only a single concentration per sample. Dilutions can be considered a form of technical replicate and provide additional information about confidence in the RFI measurement. In contrast, Western blots contain size information about the analytes being measured, which is not available in an RPPA measurement. Although the cost per antibody-sample is similar between Western blot and Zeptosens RPPA, the time required to generate many measurements is much shorter with RPPA. Lastly, RPPA requires much less sample than Western blot, allowing for many more measurements with the same sample.

In summary, compared to Western blots, the advantages of RPPA are: higher throughput and sensitivity, larger dynamic range, significantly less sample required, less antibody needed, and lower cost per data point.

## Background issues with clinical tissue samples

We encountered issues with high background noise when conducting RPPA assays with certain patient tissue samples, which could be traced to two primary causes: (1) blood contamination, and (2) autofluorescence signals innate to the tissue itself. When performing RPPA assays with patient samples, two control arrays are required to mitigate problems associated with background signals: (i) a control array stained with secondary antibody only (no primary antibody) to assess non-specific binding of the secondary antibody, and (ii) a control array with no primary or secondary antibodies to assess autofluorescence.

### Blood contamination

Human immunoglobulins, such as IgGs, present in the blood can cross react with secondary antibodies. One way to mitigate this background binding is to use secondary antibodies that have been depleted of cross-species reactive antibodies. Several vendors (e.g., Cell Signaling Technology and Abcam) sell these “pre-absorbed” secondary antibodies that increase specificity towards species-specific primary antibodies.

To estimate and normalize sample loading amounts, usually we use total protein concentrations. However, total protein concentration measurements from samples that contain blood do not differentiate between cellular protein and protein from the blood itself. It is tricky to quantitate total protein exclusive of blood protein when blood is present. Possible solutions include the use of Protein A beads to remove blood IgGs or using housekeeping proteins or ssDNA levels^15^. Note that although these normalization methods help provide a better estimate for the overall protein contribution for non-blood cells, they do not remove the contribution of any individual protein from the blood cells, which may confound any single protein or phospho-protein measurement.

### Noise from autofluorescence

In our RPPA experiments, autofluorescence background noise was observed in patient kidney tissue samples and pancreatic laser capture microdissection (LCM) samples (data not shown). Autofluorescence is a naturally occurring phenomenon that is observed in many plant and animal tissues, and is likely due to substances such as NADPH, flavins, lipofuscin, collagen, and elastin that exist in tissue and blood^16^. Other researchers have reported that the use of near-infrared fluorescence dyes and detection systems can at least partially alleviate background due to autofluorescence^1^.

**Table 1.** Background issues in different samples we tested.

| Samples | Background issues | Reasons |
| --- | --- | --- |
| cell lines | no | - |
| human pancreatic organoids | no | - |
| human pancreatic LCM samples | yes | autofluorescence |
| human kidney tumor core needle biopsies | yes | blood contamination, autofluorescence |

## Experimental case studies

### Cell line samples

We measured several total and phospho-protein levels in 15 cancer cell lines grown in standard cell culture conditions (see Data reproducibility section above). Cells were cultured in 75 cm^2^ flasks and harvested at approximately 80% confluence. All sample concentrations were adjusted to 2 mg/mL before further dilutions for chip spotting. A mixture of lysate from the 15 cell lines was prepared as a standard control and spotted onto each array to normalize between arrays. A panel of 17 antibodies was tested on all cell lines. The Zeptosens RPPA results are consistent with those of conventional Western blots carried out on the same cell lysates as shown in Figure 5, and what we expect based on the genomic backgrounds of the cell lines and consensus signaling pathway information (Figure 6). For example, PTEN signal was low in the PTEN null cells including LNCaP, LNCaP-AR, SkMel-133, and SkMel-207. A correlation between phospho-MEK1/2 and phospho-ERK1/2 levels and an inverse correlation between PTEN and phospho-AKT were observed in most cell lines, as expected.

**Figure 6.**
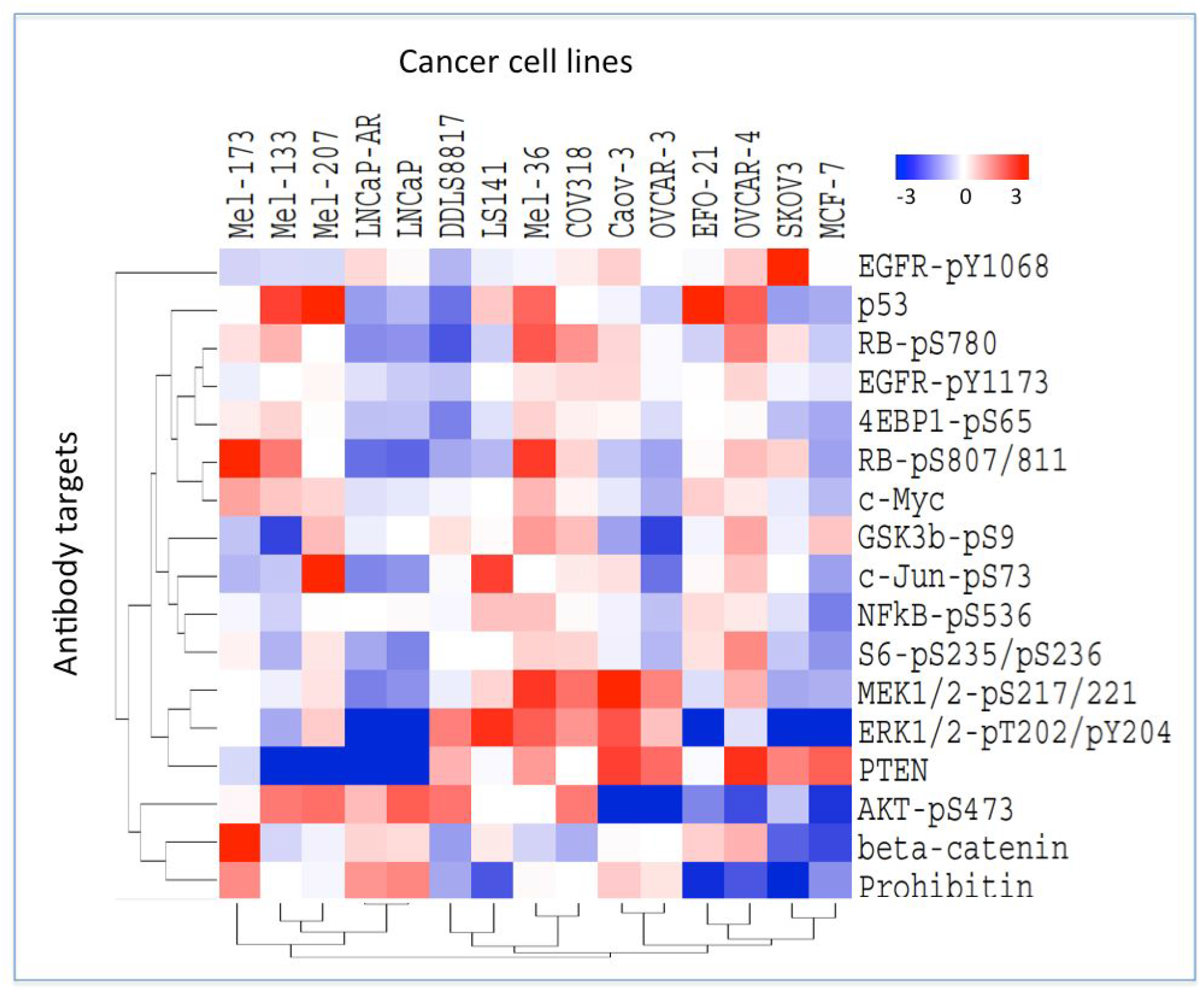
Protein profiling of cancer cell lines. Average-linkage hierarchical clustering of an uncentered Pearson correlation similarity matrix was executed with Cluster 3.0 and visualized with TreeView^17^. From the normalized RFI values, data were log transformed and median centered to generate the relative values that are visualized. See Supplemental Data 3 for all raw values.

### Tissue samples

We analyzed the spatial heterogeneity of protein concentrations in four patient kidney tumors. For each patient, core needle biopsies (20-gauge) from two or three sites across normal and tumor tissue were collected. The tissue weight of each sample was approximately 1-2 mg. The Covaris t-PREP impactor system was used to pulverize the samples in a frozen state throughout the process. A volume of 50 μL of CLB1 buffer was immediately added to the pulverized tissue. We followed the standard Zeptosens protein lysis protocol, which yielded approximately 40 μL protein lysate at a concentration of ~3-4 mg/mL. A panel of 35 antibodies was used for RPPA analysis. Some total protein and phospho-protein levels were different between tumor and normal kidney samples, and these differences were relatively consistent across the sites. For example, one antibody that showed consistent signal differences was AMPK alpha-pT172 (Figure 7).

**Figure 7.**
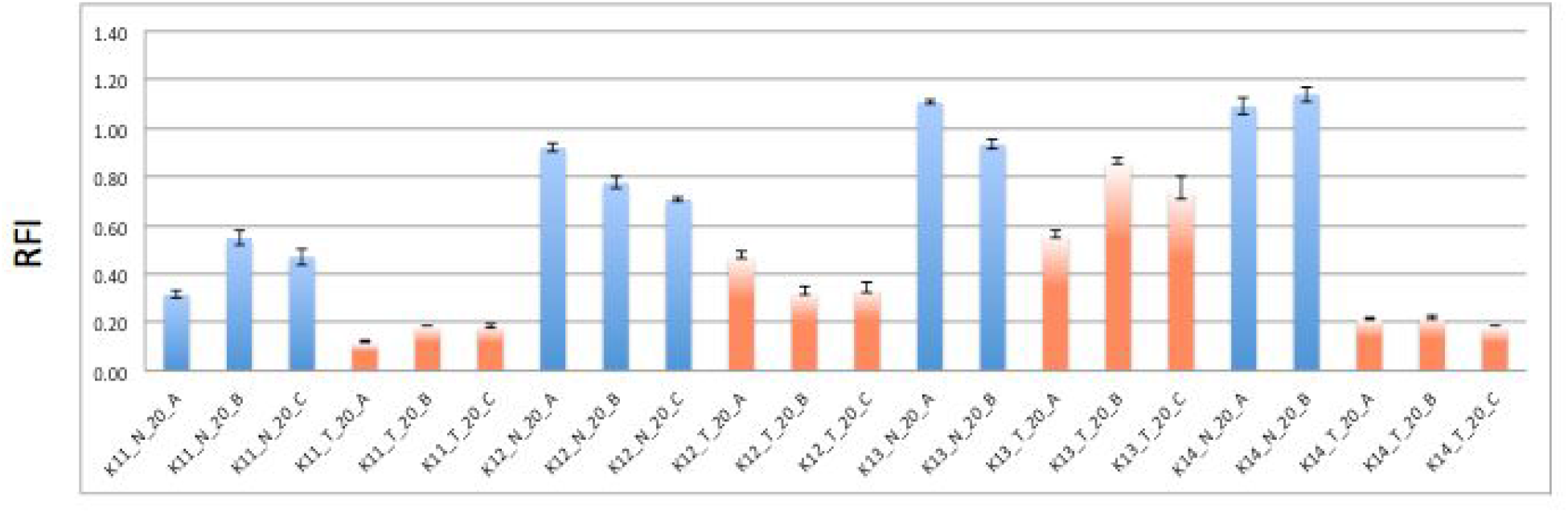
Heterogeneity in patient kidney tumors vs. normal tissues. Normalized RFI values of AMPK alpha-pT172 detected in kidney tumors from four patients (K11, K12, K13, and K14). Multiple core needle biopsies from several sites were evaluated (A, B, and C). N = normal (blue) and T= tumor (orange). The error bars are from three technical replicates on the same array.

## Advantages and disadvantages of the RPPA technology

RPPA is a powerful tool to quantitatively measure hundreds of signaling proteins in cultured cells and clinical samples. The power of this technology lies in the ability to simultaneously probe hundreds of samples with many specific antibodies, which provides sensitive quantification of proteomic expression and post-translational modification status. By requiring minute amounts of material, it is well suited for probing signal transduction in a variety of sample types including cultured cells, organoids, and patient tissues, for example, core needle biopsies, LCM samples, and formalin fixed and paraffin embedded (FFPE) tissues^1,18^. The primary challenge with RPPA technologies is the requirement for highly specific antibodies as no secondary size selection is utilized (Western blot), no sequencing by M/Z profile occurs (mass spectrometry), and it does not take advantage of multiple antibodies to increase specificity (ELISA). However, compared with other commonly used protein analysis platforms such as Western blot, ELISA, immunohistochemistry, and mass spectrometry, RPPA exhibits robust quantification, requires small amounts of sample due to high sensitivity, allows for high-throughput processing of samples, and enables measurements of total protein as well as phosopho-proteins and other post-translational modifications. RPPA represents a promising and sensitive tool to identify new biomarkers and potential therapeutic targets.

## Acknowledgments

We thank the Beene Core Facility at MSKCC for assistance, the MSKCC Department of Pathology for support, members of the Sander Group Pi-Club for ongoing discussions and collaboration, and Laura Kleiman for feedback on the manuscript. We thank Michael Pawlak and his lab in Tübingen for valuable advice regarding optimization of our in-house Zeptosens platform, and staff at the Zeptosens and GeSim commercial providers for technical assistance.

We are grateful to Gordon Mills and his staff at MD Anderson for introducing us to their RPPA technology several years ago, for extensive collaboration, as well as for expert advice. We thank James Hsieh for providing kidney tissue, and Steven Leach for the pancreatic LCM samples and organoids.

